# transomics2cytoscape: An automated software for interpretable 2.5-dimensional visualization of trans-omic networks

**DOI:** 10.1101/2023.03.08.531686

**Authors:** Kozo Nishida, Junichi Maruyama, Kazunari Kaizu, Koichi Takahashi, Katsuyuki Yugi

## Abstract

Biochemical network visualization is one of the essential technologies for mechanistic interpretation of omics data. In particular, recent advances in multi-omics measurement and analysis require the development of visualization methods that encompass multiple omics data. Visualization in 2.5 dimension (2.5D visualization), which is an isometric view of stacked X-Y planes, is a convenient way to interpret multi-omics/trans-omics data in the context of the conventional layouts of biochemical networks drawn on each of the stacked omics layers. However, 2.5D visualization of trans-omics networks is a state-of-the-art method that primarily relies on time-consuming human efforts involving manual drawing. Here, we present an R Bioconductor package ‘transomics2cytoscape’ for automated visualization of 2.5D trans-omics networks. We confirmed that transomics2cytoscape could be used for rapid visualization of trans-omics networks presented in published papers within a few minutes. Transomics2cytoscape allows for frequent update/redrawing of trans-omics networks in line with the progress in multi-omics/trans-omics data analysis, thereby enabling network-based interpretation of multi-omics data at each research step. The transomics2cytoscape source code is available at https://github.com/ecell/transomics2cytoscape.

## Introduction

Visualization of large-scale molecular networks and comprehensive data is an essential technology to facilitate the interpretation of ‘omic’ data^1^. To date, a variety of network visualization software has been developed such as VANTED^2^ for metabolomic data in the context of metabolic pathways and Cytoscape^3^ and OmicsNet^4,5^ for protein-protein interactions and other types of networks.

Yugi and colleagues have established a ‘trans-omics’ analysis method that allows the reconstruction and visualization of biochemical networks spanning multiple omic layers^6^. Their principle for visualizing the trans-omic network is to provide interpretability with the support of researchers’ prior knowledge. The prior knowledge used to visualize interpretable networks has two components: (1) conventional pathway map layout and (2) stacked omic layers with a 2.5D perspective, which is an isometric view of the stacked layers in a 3D space. However, the visualization of trans-omic networks was dependent on personal and time-consuming efforts, especially in the case of preparing a submission-ready quality figure. Accordingly, it was not realistic to visualize trans-omic networks at each step of a multi-omics/trans-omics research project, which would allow one to refer to a preliminary image of the network for the further characterization of molecular mechanisms in subsequent steps.

Here we present an R package, transomics2cytoscape, that automates the visualization of trans-omic networks from tabular data describing network connectivity. The transomics2cytoscape package reproducibly generates images of the trans-omic networks in the context of conventional pathway map layouts and stacked omic layers. In line with recent developments in workflow automation, transomics2cytoscape allows one to bypass human-dependent processes in the visualization of trans-omic networks. In this article, we present an overview of transomics2cytoscape along with two use cases visualizing previously published trans-omic networks.

## Results

### Software overview

We developed the R package transomics2cytoscape that automates the 2.5D visualization^7,8^ workflow for trans-omics studies. transomics2cytoscape has two types of input table files and two functions that process and visualize each of the input table files. An understanding of these file formats and how to prepare the data is required to use transomics2cytoscape. One of the two input table files is called the “layer definition table file”, which specifies the pathway network of each layer and its Z-coordinate, and the other file is called the “trans-omic interaction table file”, which defines the interaction that connects each layer (Fig. 1, top). Hereafter, we refer to an interaction between omic layers as a “trans-omic interaction”. Given the layer definition file as an input, create3Dnetwork draws multiple layers in 3D space.

**Fig. 1:**
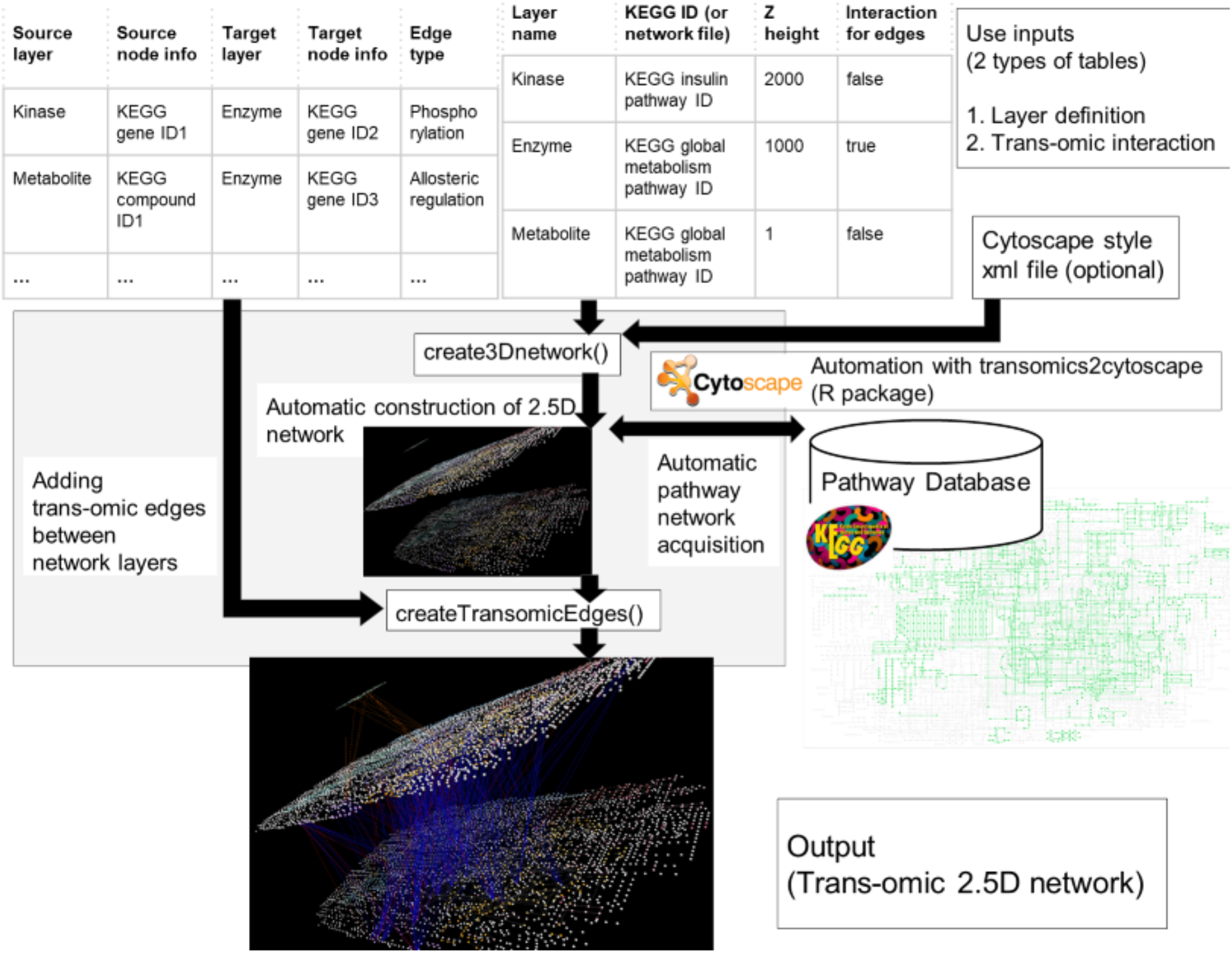
The transomics2cytoscape visualization workflow. The user is expected to prepare table files in two formats. One is an input for the “create3Dnetwork” function to create multiple omic layers and the other is an input for the “createTransomicEdges” function to create edges for the trans-omic interactions drawn by create3Dnetwork. The “create3Dnetwork” function can automatically import network layouts from the KEGG database by entering the pathway ID information in the table. The user can change the default style by preparing an XML file for a different Cytoscape style.

The data format of the layer definition table file is a four-column tab-delimited text file (Fig. 2a). The first column is for an arbitrary character string representing an index for each layer. This information is also used in the trans-omic interaction table file. The second column is expected to contain a KEGG pathway ID or a filename in the file format that Cytoscape can import. For KEGG, there is no need to prepare file data; just write the KEGG pathway ID and transomics2cytoscape will retrieve the pathway data from KEGG. If a user needs to import original network data other than KEGG, the user should place a Cytoscape-compatible file in the networkDataDir path, which is one of the arguments of the create3Dnetwork function. The third column is the position in the Z-coordinate of each network layer. The fourth column is a Boolean value indicating whether or not trans-omic interactions should be provided with the edges as well as the nodes. Trans-omic interactions apply not only to edges from a molecule (node) in a layer to a molecule (node), but also to edges from a molecule (node) in a layer to an interaction (edge) in a layer. For example, the global metabolic pathway map of KEGG (https://www.genome.jp/pathway/map01100) does not have nodes to represent molecular interactions (i.e. metabolic reactions). By setting the fourth column to “true”, transomics2cytoscape adds a node at the midpoint of a network edge in the layer. By changing the network layout, the user can also visualize the node-to-edge (and the opposite) and edge-to-edge interactions, which cannot be done with the original KEGG network layouts. The number of layers in this table file can be increased to any number. Fig, 2a shows examples of trans-omic networks with three layers (a-1), and five layers (a-2).

**Fig. 2:**
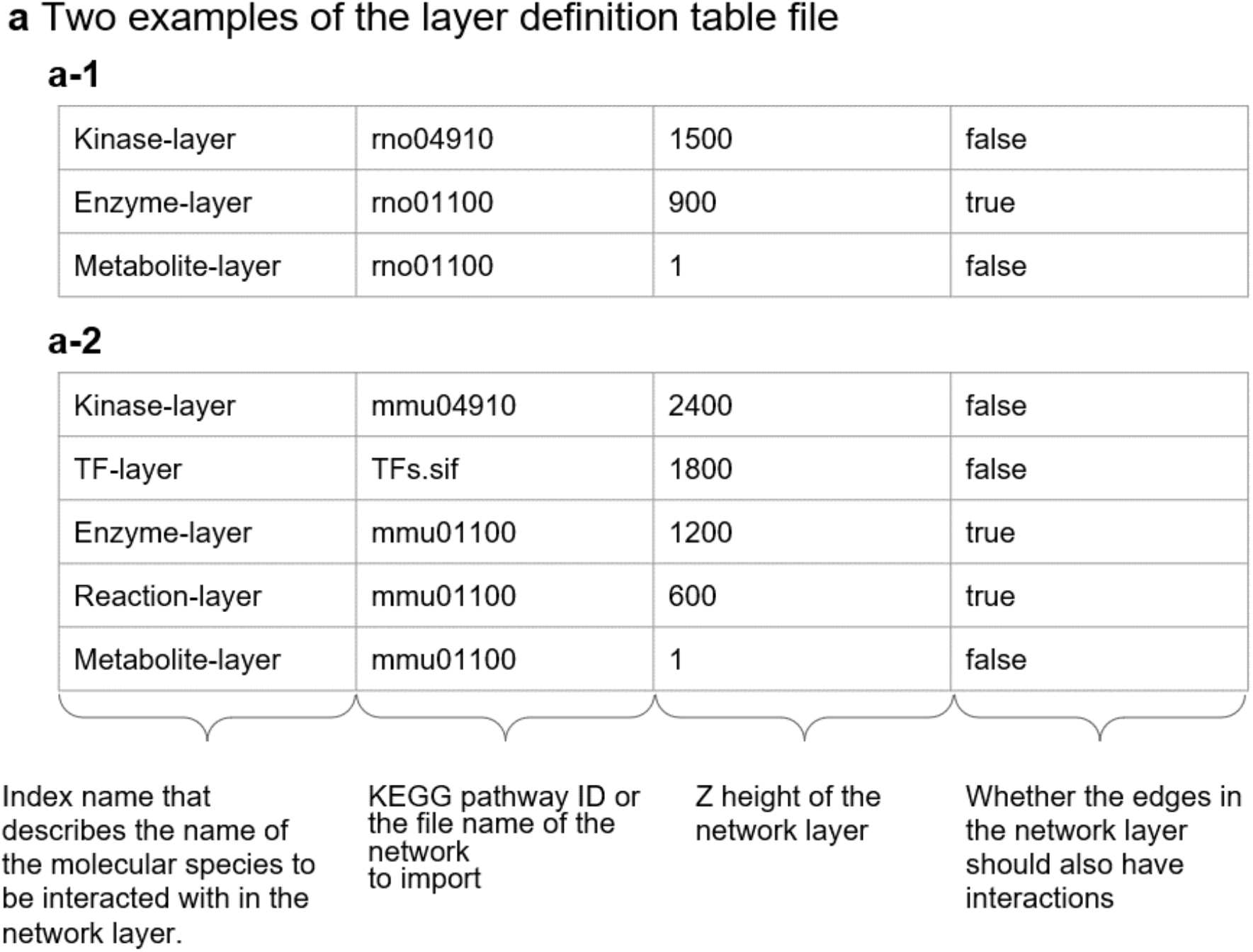
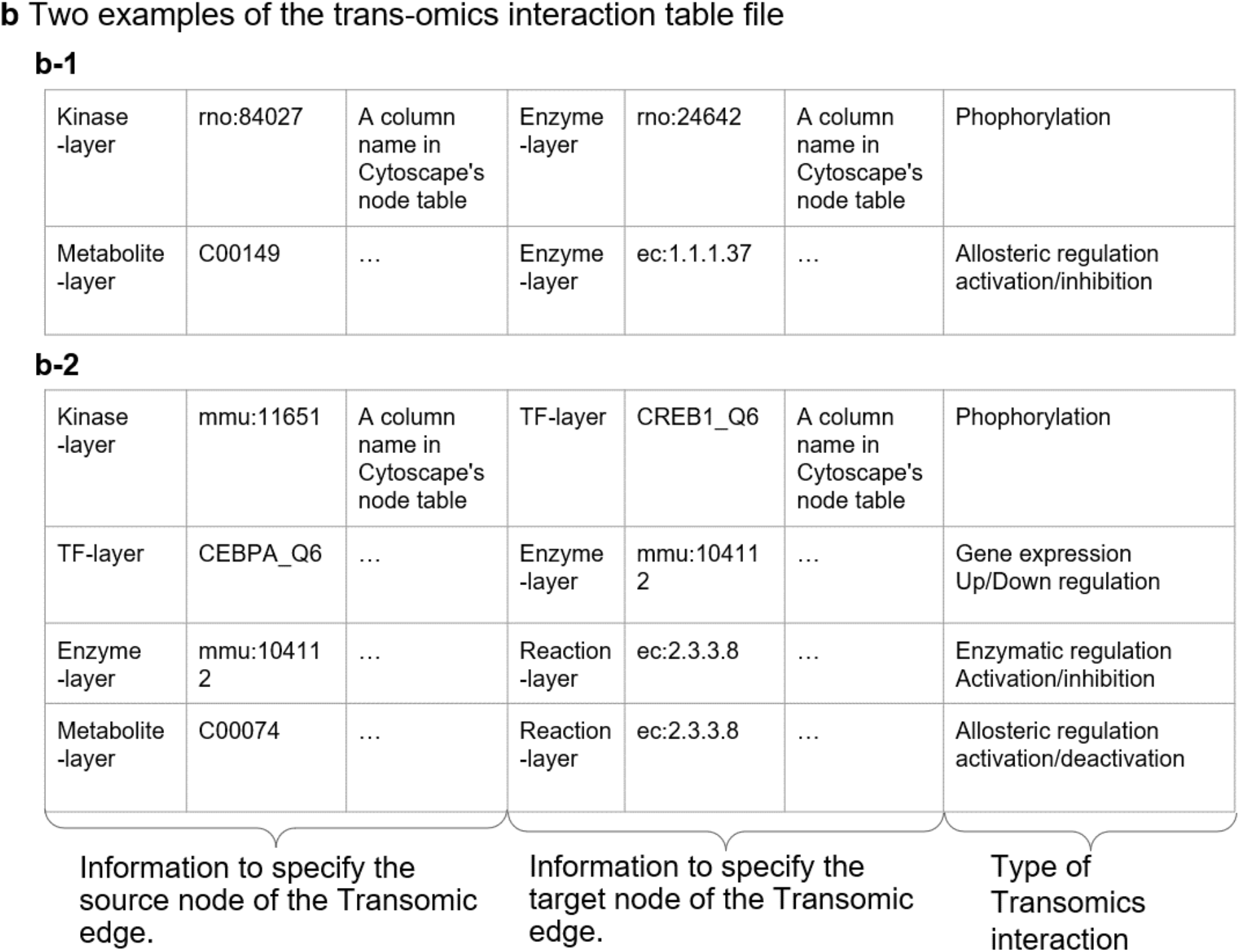
Description of input table formats of transomics2cytoscape. (a) Two examples of the layer definition file. The first column is for an index for each layer. The second column is for a KEGG pathway ID or a filename that Cytoscape can import. The third column is the position of each network layer in the Z-coordinate. The fourth column indicates whether a node should be created at the center of an edge. In this example, we provide molecule types as the layer index. (b) Two examples of a trans-omic interaction table file. In the first three columns, transomics2cytoscape requires information specifying the source nodes of the trans-omic interaction. The next three columns are used for information specifying the target nodes of the trans-omic interaction. The seventh column describes the type of trans-omic interaction.

The trans-omic interaction table is a tab-delimited text file with seven or more columns (Fig. 2b). The first to third columns and the fourth to sixth columns contain information about the source (start) and the target (goal) of the trans-omic interaction, respectively. The first and fourth columns are character strings corresponding to the first column of the layer definition file: the source layer and the target layer of the trans-omic interaction, respectively. The second column is for designating the ID of the source node of the trans-omics interaction. The third column specifies the column name in the node table of Cytoscape. The fifth and sixth columns are the same as the second and third columns except that the fifth and sixth columns are for the target node of the trans-omics interaction. The seventh column is for an arbitrary character string that describes the attribute value of the interaction between layers (e.g. “metabolite_activator” or “metabolite_inhibitor”). Fig. 2b-1 and b-2 are the trans-omic interaction table files for the layer definition table files in Fig. a-1 and a-2, respectively.

Molecular species in the same layer may belong to different ID namespaces depending on the contexts in which the entities are specified, for example, as an enzyme protein or an enzymatic reaction. The namespace of the target IDs for metabolite-protein interactions, which spans from the metabolite layer to the enzyme layer, is the EC number (Fig. 2b-1), because the BRENDA^9^ database, a major data source for metabolite-protein interactions, specifies each enzyme reaction by an EC number. The namespace of the target IDs of protein phosphorylation, which spans from the kinase layer to the enzyme layer, is the ID of the enzyme protein, which is typically provided after proteomic measurements (e.g., UniProt^10^). Also, the KEGG global metabolic pathway map, which transomics2cytoscape expects to be one of the most frequently used data sources for a network layer, does not use EC numbers but KEGG reaction IDs for metabolic enzymes. transomics2cytoscape converts EC numbers to KEGG reaction IDs using the ec2reaction function, so that it can connect trans-omic interactions of different modalities that meet at the metabolic enzyme layer (Fig. 3).

**Fig. 3:**
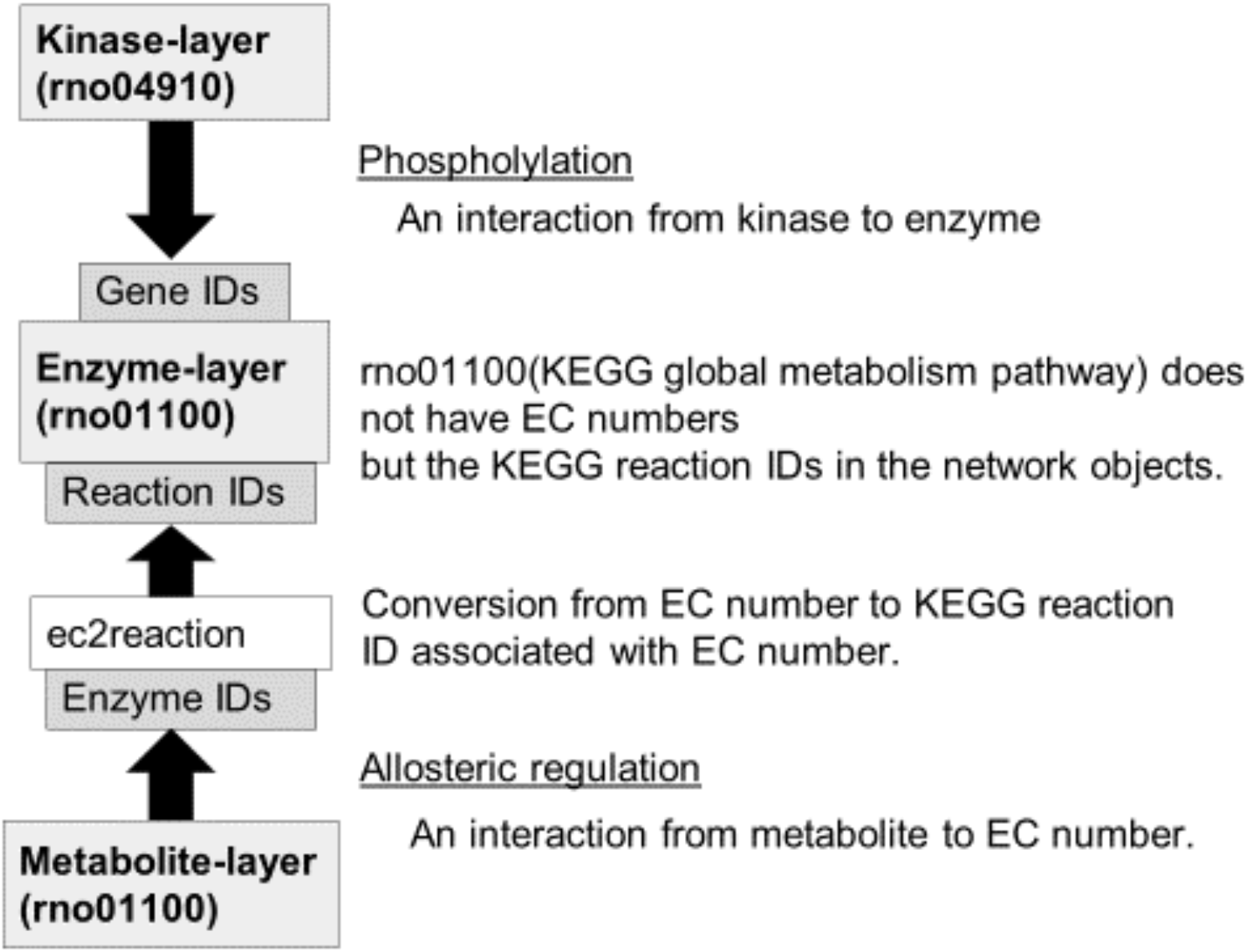
Bridging multiple ID namespaces for an identical enzyme targeted by different types of trans-omic interactions. The ec2reaction function converts an EC number into a KEGG reaction ID to integrate trans-omic interactions that have different IDs for the same enzyme. For example, the target ID for regulatory metabolite-protein interactions available in BRENDA is an EC number, whereas the target ID for phosphorylation of metabolic enzymes is typically a gene ID. Conversion of IDs, such as from an EC number to a KEGG reaction ID, allows connecting these trans-omic interactions to the metabolic enzymes that they target.

### Case study I: Insulin action across the phosphoproteome and metabolome in rat hepatoma Fao cells

First, we show an example of visualizing a trans-omic network for acute insulin action in rat hepatoma Fao cells^6^. This network does not include gene expression layers but includes metabolic regulatory pathways that traverse the phosphoproteome and metabolome layers. transomics2cytoscape provides an automated workflow that facilitates creating a 2.5D network from the table data. We created the trans-omic network of insulin action (Fig. 4) using datasets in the folder named ‘usecase1’ at https://doi.org/10.5281/zenodo.7471290 and codes in the folder named vignettes at https://doi.org/10.5281/zenodo.7471290.

**Fig. 4:**
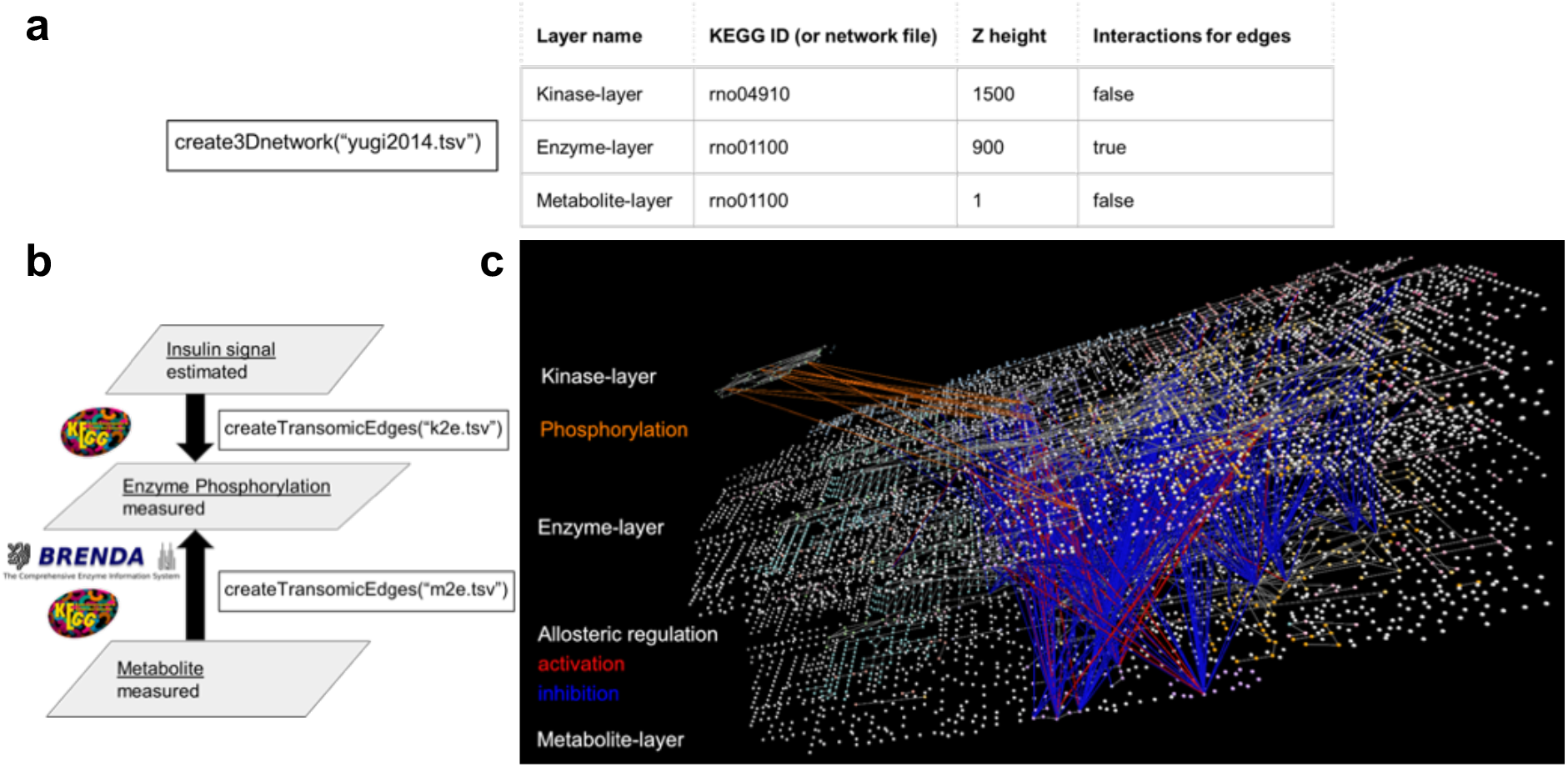
A three-layered trans-omic network of insulin action in rat hepatoma Fao cells automatically generated by transomics2cytoscape. (a) A layer definition table file (‘yugi2014.tsv’) that defines names, positions (‘Z height’), etc. of the three layers. (b) The arguments to the createTransomicEdges function, such as ‘k2e.tsv’ and ‘m2e.tsv’, are names for trans-omic interaction table files expected to contain kinase-substrate relationships and metabolite-protein interactions, respectively. (c) The trans-omic network of insulin action in rat hepatoma Fao cells^6^ automatically produced by transomics2cytoscape. The top layer is the insulin signaling pathway imported from KEGG (rno04910). The middle and bottom layers are the global metabolic pathways from KEGG (rno01100) for enzymes and metabolites, respectively. The nodes in the network are defined in the same way as in KEGG. Spheres represent metabolites and other molecules and cubes represent gene products, typically proteins. Additional nodes representing enzymes were drawn at the midpoint of all reaction edges of the global metabolic pathway map (rno01100) since it does not contain any nodes for enzymes, but only edges. As for the edge color between layers, orange indicates protein phosphorylation. Red and blue indicate activation and inhibition of an enzyme by metabolite-protein interaction, respectively.

### Case study II: Responses to oral glucose challenge across protein phosphorylation, transcriptome and metabolome layers in the liver of wild type and obese mice

Our second example is a trans-omic network in the wild type (WT) and obese (*ob*/*ob*) mouse liver responding to oral glucose challenge^11^. This network includes gene expression layers that are not present in the Case study I network. transomics2cytoscape can easily expand the number of layers in its visualization. We show an automatic visualization of a trans-omic network that integrates protein phosphorylation, transcriptome and metabolome data obtained from the liver of WT and *ob*/*ob* mice.

The network has trans-omics interactions between five layers i.e. from signaling molecules to transcription factors (TFs), from TFs to enzymes, from enzymes to reactions, and from metabolites to metabolic reactions. The five layers of the network are the KEGG Insulin signaling pathway (mmu04910), a layer for TFs (tfs.sif file), and the KEGG global metabolic pathway (mmu01100) for the remaining three layers (enzymes, reactions, and metabolites). We created the trans-omic network of the glucose response in the mouse liver (Fig. 5) using datasets in the folder named usecase2 at https://doi.org/10.5281/zenodo.7471290 and codes in the folder named vignettes at https://doi.org/10.5281/zenodo.7471290.

**Fig. 5:**
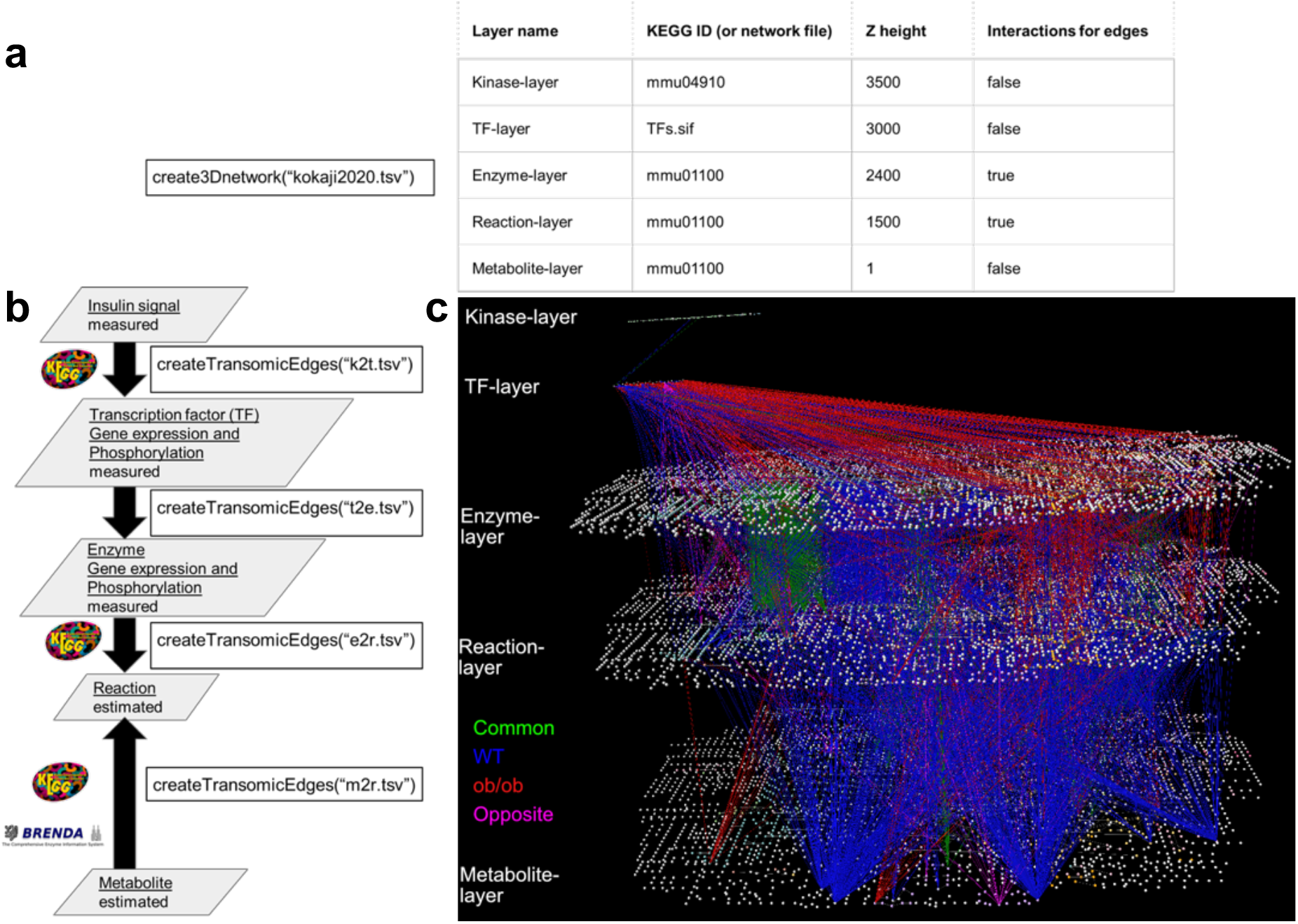
transomics2cytoscape visualization of a five-layered trans-omic network of the responses to oral glucose challenge in the mouse liver. (a) A layer definition table file (‘kokaji2020.tsv’) that defines names, positions (‘Z height’), etc. of five layers. Note that the layers are functional classes of molecules and do not necessarily correspond to omic layers. (b) The arguments for the function createTransomicEdges such as ‘k2t.tsv’, ‘t2e.tsv’, ‘e2r.tsv’, and ‘m2r.tsv’ are names for trans-omic interaction table files expected to include kinase-substrate (TF) relationships, TF-enzyme (target gene) relationships, enzyme-reaction relationships, and metabolite-reaction (enzyme) interactions, respectively. (c) The trans-omic network of the responses to glucose challenge in the liver of WT and *ob*/*ob* mice^11^ automatically produced by transomics2cytoscape. The top layer is the insulin signaling pathway imported from KEGG (mmu04910). The second layer from the top is for transcription factors. The third to fifth layers from the top are the global metabolic pathways of mice from KEGG (mmu01100), representing enzyme genes, metabolic reactions and metabolites, respectively. The edges between each layer are trans-omics interactions from signaling molecules to transcription factors (between layers 1 and 2), from TFs to target enzyme genes (between layers 2 and 3), from enzyme genes to metabolic reactions (between layers 3 and 4), and from metabolites to metabolic reactions (between layers 5 and 4). The edges in blue, red, green, and magenta indicate regulation functioning: only in WT, only in *ob*/*ob*, in similar ways both in WT and *ob*/*ob*, and in opposite ways in WT and *ob*/*ob*, respectively.

### Comparing transomics2cytoscape with existing tools

There are four specifications required for software visualizing a trans-omic network:

1.Importing pathway map files provided by pathway databases [e.g. KGML files of KEGG^12^]

2.Stacking 2D pathways as layers of a 2.5D network

3.Drawing edges between stacked layers

4.Automated workflow

Compatibility with pathway resources provided by KEGG is indispensable because the KEGG PATHWAY is one of the most commonly used and referred resources among various pathway databases. VANTED-HIVE^13^ was employed in the trans-omic network visualization by Yugi and colleagues^6^. The specification ‘1’ shown above is provided by its original software VANTED, and ‘2’ and ‘3’ are realized in its extension VANTED-HIVE. Other software such as Arena3D^14^ and BioLayout Express 3D^15^ were also considered for this purpose by the authors of Ref. 6 but were not satisfactory in some aspects (Table 1).

**Table 1.** Comparison of transomics2cytoscape with existing tools. Unique to transomics2cytoscape is that it allows automatic visualization of biochemical networks in a stacked-layer manner after importing conventional pathway layouts.

|  | transomics2cytoscape | VANTED-HIVE | BioLayout Express 3D | Arena3D | OmicsNet |
| --- | --- | --- | --- | --- | --- |
| Application type of 3D visualization | Desktop Application | Desktop Application | Desktop Application | Web Application | Web Application |
| Whether the dataset is limited to specific species | Not limited to species | Not limited to species | Not limited to species | Not limited to species | One to ten species |
| Is actively maintained | ++<br>(Bioconductor package) | — | + | ++ | ++ |
| Requirements for the visualization of a trans-omic network |  |  |  |  |  |
| Importing pathway map files provided by pathway databases | + | + | — | — | — |
| Stacking 2D pathways as layers of a 2.5D network | + | + | — | + | + |
| Drawing edges between stacked layers | + | + | — | + | + |
| Automated workflow | ++ | — | — | — | + |
| 3D network visualization |  |  |  |  |  |
|  | transomics2cytoscape | VANTED-HIVE | BioLayout Express 3D | Arena3D | OmicsNet |

**Table1. Comparison of transomics2cytoscape with existing tools (cont'd)**
|  |  |  |  |  |  |
| --- | --- | --- | --- | --- | --- |
| How to stack networks as layers | Each KEGG pathway or user-defined network | Each KEGG pathway or user-defined network | Stacking is not supported | User-defined network | Pre-defined (Microbes, Metabolites, Genes/proteins, SNPs) |
| Layout of networks in layers | Conventional layout of biochemical networks based on KEGG pathways | Conventional layout of biochemical networks based on KEGG pathways | — | Automatic layout by network analysis algorithm | Automatic layout by network analysis algorithm |
| Automation of the data integration and visualization workflow |  |  |  |  |  |
| Automation target | Trans-omics | No workflow automation | No workflow automation | No workflow automation | Multi-omics |
| Automation customizability | It works locally and is scriptable in R. | No workflow automation | No workflow automation | No workflow automation | The web application part is not open source and cannot be customized. |
| Data integration target | KEGG pathways and arbitrary molecular interactions | Arbitrary molecular interactions and biological images | Arbitrary molecular interactions | Arbitrary molecular interactions | Protein-protein, miRNA-gene, Metabolite-protein, TF-gene in limited species |
| Input data of the automated workflow | List of network files supported by Cytoscape (or KEGG ID), List of Transomic interactions | No workflow automation | No workflow automation | No workflow automation | List of molecule IDs |

One limitation of VANTED-HIVE is that it does not support specification ‘4’, which is automation of the workflow for visualizing trans-omic networks. Visualization of the trans-omic network in Ref. 6 demanded a considerable amount of manual effort and was dependent on the users’ personal skills and patience. In addition, VANTED-HIVE does not appear to be maintained anymore. OmicsNet is a web platform for multi-omics integration and 3D network visualization. It can automatically execute workflows from data integration of multi-omics data to 3D network visualization. However, the data sources that are predefined in OmicsNet are limited to certain species. Furthermore, the users cannot customize the data integration workflow. For example, the users cannot define layers in the 2.5D network other than the predefined settings that are designed on the basis of data object classification (e.g. microbes, metabolites, genes/proteins, and SNPs), though there are users that need to define the layers in other ways, such as the functional/mechanistic classifications shown in Case studies I and II (e.g. signaling molecules, TFs, target genes, metabolic reactions etc.). In addition, the layout of the network on each layer is limited to automatic coordination. The users cannot import conventional biochemical pathway layouts such as those in the KEGG PATHWAY database.

### Prior knowledge is necessary for creating an interpretable network

People are likely to perceive, recognize and interpret networks dependent on their prior knowledge, which is implemented as conventional layouts of the networks. Do readers immediately find what the networks in Figs. 6a and b represent? These networks are the London underground network inside the Circle Line redrawn based on algorithms named degree sorted circle layout and compound spring embedder (CoSE^16^), respectively. Residents in London are expected to recognize their present location, destination and the transfer stations in reference to the Circle Line in the original conventional layout (Fig. 6c). However, it is difficult to find where the Circle Line is in the algorithmically optimized network layouts.

**Fig. 6:**
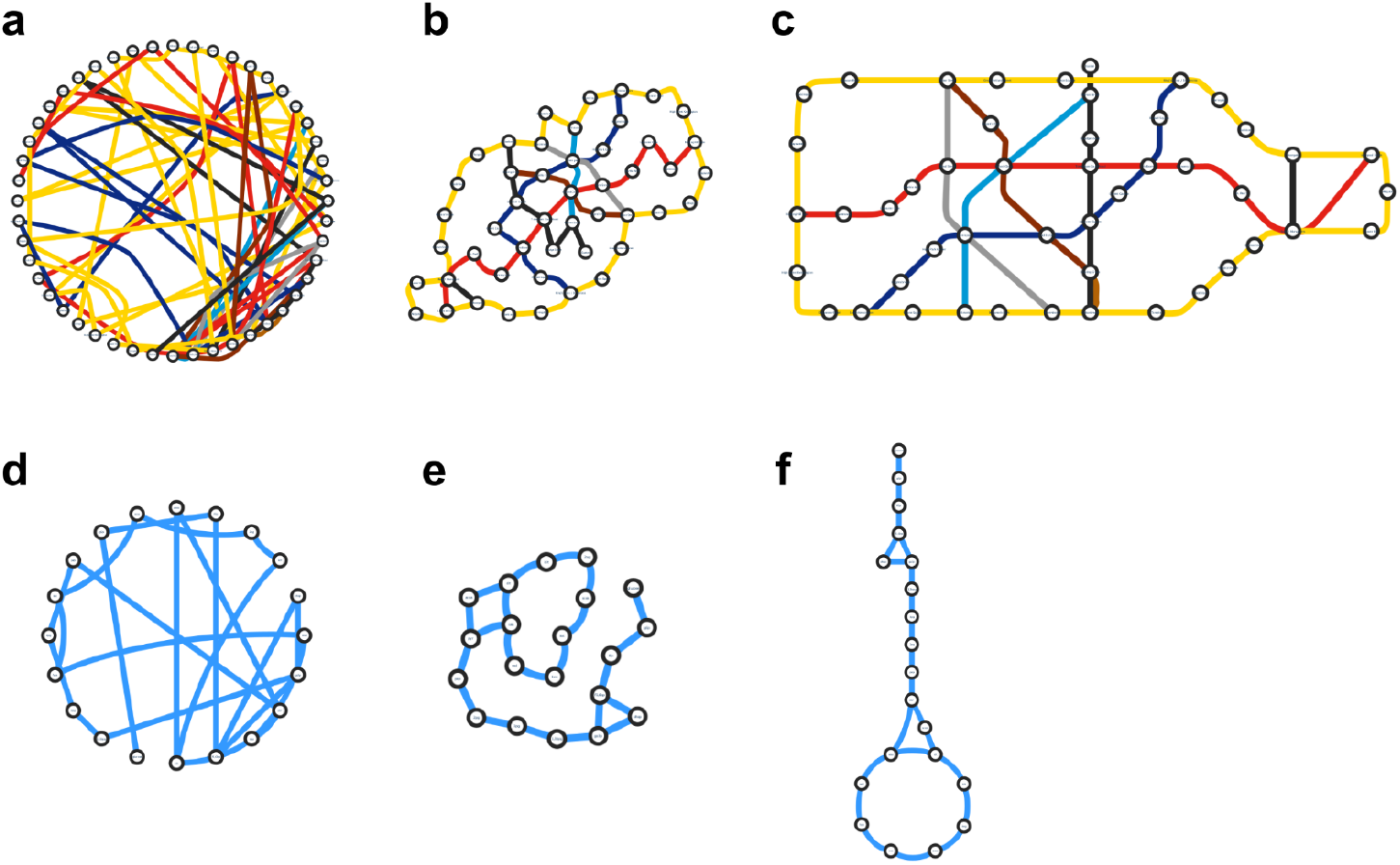
Unbiased network layouts are not necessarily more easily interpretable than conventional layouts. **(a**-**c**) The London underground network inside Circle Line before 2009 in three different layouts. The network layouts **a** and **b** are generated from the conventional layout **c** based on degree sorted circle layout and compound spring embedder (CoSE^16^), respectively. (**d**-**f**) The glycolysis and the tricarboxylic acid cycle in three layouts as in **a**-**c**. These images show that the auto-layout algorithms in common use produce layouts that differ significantly from the conventional pathway layouts.

Prior knowledge is similarly essential in recognition of biological networks. The networks in Figs. 6d and e depict central carbon metabolism redrawn by degree sorted circle layout and CoSE, respectively. In the conventional pathway layout (Fig. 6f), biologists working on metabolism might recognize the positions of the metabolites and the enzymes of interest in reference to the TCA cycle in the lower part of Fig. 6f, though it is difficult with the distorted TCA cycle in the algorithmically generated network layouts, which are not distinguishable from the underground network (Compare Figs. 6a and d).

If a network layout is shared as prior knowledge in a certain community, the network should be visualized according to the shared prior knowledge but not in a hairball layout or another algorithmically generated layout. KEGG pathway maps are used as such shared prior knowledge, particularly in the metabolism field. Import of the KEGG layouts to software facilitates the prior knowledge-based visualization and interpretation of networks.

## Discussion

The transomics2cytoscape package enables automatic visualization of 2.5D networks ^6,11^ in conventional pathway layouts. Using the package, it is now possible to draw trans-omic networks at any step of a multi-omics/trans-omics research project, thereby allowing access to preliminary networks at each step. This information may provide hints to explore novel molecular mechanisms across multiple omic layers.

The transomics2cytoscape package automates only the 2.5D visualization part of trans-omics analysis, but not other processes such as omic data integration. Automation of the workflow of trans-omics analysis before and after visualization, which is described below, will be presented in our future works. We plan to provide a workflow execution environment that summarizes all of these processes.

The connectivity data of trans-omic networks are provided after reconstruction procedures, but these are not fully automated yet. Fully automated network reconstruction and connecting with transomics2cytoscape will provide a seamless end-to-end pipeline from omic data to visualization. We have established a workflow of network reconstruction that includes identifying changed molecules from multiple omic data and connecting the changed molecules based on prior knowledge, such as publicly available pathway databases. Automation of the procedure has already been prototyped in our group using the Garuda Platform^17^, which allows cooperative execution of multiple software packages that constitute each step of the end-to-end pipeline. We plan to define file formats that facilitate data exchange and cooperative procedures with the Garuda-compatible pathway reconstruction software.

A reconstructed 2.5D network is also an information-rich representation from which to extract biological relevance. Kokaji *et al*.^11^ developed a technology to summarize 2.5D networks by using enrichment analysis for congregating a group of molecules into a name of a pathway. For example, in a case where the fraction of dynamically expressed genes in the glycolysis pathway is significantly larger than in other pathways, and the genes encoding the enzymes are regulated by a common transcription factor X, it can be summarized “enzyme expression in glycolysis is significantly changed by regulation under X”. We will consider implementing this summarizing function in the future.

Currently, transomics2cytoscape supports only the KEGG ID and pathway file format. In the future, we will extend this to other BioPAX^18^-compatible pathway databases such as Reactome^19^ and Wikipathways^20^. An alternative format is Systems Biology Graphical Notation (SBGN), which provides a universal notation rule for intracellular biochemical processes such as metabolism, gene expression regulation, signal transduction, and other regulatory pathways^21^. Large-scale pathway maps such as EGF signaling by Oda *et al*.^22^ and the Parkinson’s disease-related pathway by Fujita *et al*.^23^ have already been published in the SBGN format.

We are considering future addition of functions that support generating 3D movies of trans-omic networks, such as previously presented in the study of insulin action in rat hepatoma Fao cells^6^, in which one of the present authors participated. The movie first shows the perspective of the trans-omic network and then shows close-ups of each pathway and molecule so that viewers can be aware of the connection between the entire network and each component. The movie was authored using a 3DCG software Shade (https://shade3d.jp). However, the authoring process was labor-intensive and required expertise specific to the software. Further automation will be needed to make the movie-authoring technology accessible to a broader usership. CyAnimator, a plug-in of Cytoscape, can automate pathway video creation^24^. However, it only supports pathways on a 2D plane and the development community is not currently very active. The BioLayout Express 3D mentioned above also has the ability to create animations. Example movies that were made six years ago are available on YouTube (https://www.youtube.com/channel/UCLUnA39yq1C0pmRNU40NRHQ). Another possibility in this area is future extension of Cy3D^25^, which is used inside transomics2cytoscape.

Although it would be another work entirely after implementing 2.5D network movie generator, it will be the next challenge for large-scale pathway visualization to provide virtual reality (VR) experiences in the trans-omics network for knowledge discovery. The VRNetzer platform is a software that enables users to experience large-scale molecular networks via VR headsets^26^. Cy2VR (https://github.com/larsjuhljensen/Cy2VR) is a tool to convert a Cytoscape network into a VR experience. In addition, a paper on visualization methods for interpreting genome-wide association study data in VR has already been published^27^. VR for visualization of single-cell data has also been presented^28^. It is worth keeping an eye on the trend of developments regarding what humans can find by wearing headsets and moving around in a large-scale network and related omics data.

## Methods

### Software components used in transomics2cytoscape

transomics2cytoscape uses KEGGREST to retrieve KGML (KEGG pathway XML). The transomics2cytoscape package in R uses RCy3^29^ to automate^30^ importing multiple pathway maps and integrating them into a 2.5D network in Cytoscape. Also, the Cytoscape Apps KEGGscape^31^ and Cy3D are used for KEGG pathway import and 2.5D network visualization, respectively.

### Dataset preprocessing

The connectivity data of the trans-omic networks shown in Yugi *et al*.^6^ and Kokaji *et al*.^11^ were obtained from supplementary datasets of the original articles, in particular Table S3 (https://www.cell.com/cms/10.1016/j.celrep.2014.07.021/attachment/5b83010d-f4f0-4c88-829f-6ceea27fd1e8/mmc4.xlsx) and Table S4 (https://www.cell.com/cms/10.1016/j.celrep.2014.07.021/attachment/833df0b2-8abc-4229-aa97-96d07f8579ba/mmc5.xlsx) of Yugi *et al*. and Data file S11 (https://www.science.org/doi/suppl/10.1126/scisignal.aaz1236/suppl_file/aaz1236_data_file_s11.xlsx) of Kokaji *et al*. The IDs of molecules were converted into KEGG–compatible IDs such as KEGG geneID and EC numbers by using Ensembl BioMart^32^ and ‘Database to Database Conversions’ provided by bioDBnet^33^ in case cross reference tables were not available.

The data used in this study are available in our GitHub repository https://github.com/ecell/transomics2cytoscape.

### Code availability

transomics2cytoscape is open source and can be found on GitHub at https://github.com/ecell/transomics2cytoscape under the Artistic-2.0 license.

## Acknowledgements

We thank Thao Tran Thanh Nguyen (Chemistry and Biomolecular Sciences, University of Ottawa) for her advice in designating the layer definition table. This work is supported by JST, CREST Grant Number JPMJCR22N5, by funding from JSPS KAKENHI grants JP18H05431 and by the Inamori Research Grants Program from the Inamori Foundation.

## Contributions

K.Y. conceived and designed the study. K.N. implemented transomics2cytoscape with help from J.M., K.K., K.T., and K.Y. K.Y. wrote the paper. K.K. reviewed the content.

